# Sex-linked Lung Estrobolome May Contribute to Pulmonary Hypertension Penetrance of *Bmpr2 R899X* Mutation via an ET-1^high^ Endoregulatory Macrophage Phenotype

**DOI:** 10.64898/2026.06.08.729693

**Authors:** O Loya, ES Villarreal, A Carneiro, S Agarwal, D Fraidenburg, J Sun, V de Jesus Perez, T Lahm, SD Oliveira

**Author notes:** **Correspondence: Suellen D. Oliveira, MS, PhD, FAHA:** Assistant Professor - University of Illinois Chicago; College of Medicine; 835 S. Wolcott Ave (M/C 513) Medical Sciences Building Room E713; Chicago, IL, USA - 60612.

## Abstract

Mutations in the bone morphogenetic protein receptor 2 (BMPR2) are a major genetic driver of pulmonary arterial hypertension (PAH), yet their penetrance is strikingly sex-biased: females are disproportionately affected, while males experience poorer outcomes. While hormonal and chromosomal factors have been implicated, the biological basis for this disparity remains not fully understood. Here, we investigated the role of the lung microbiome in sex-linked PAH pathogenesis. We hypothesized that increased BMPR2 mutation penetrance in females is partly driven by the accumulation of potent vasoactive molecules, such as endothelin-1 (ET-1), in response to lung microbiome dysbiosis. Using humanized Bmpr2^+/R899X^ mice, we integrate lung metagenomics with basic functional immune profiling to show that females develop a distinct microbiome profile, characterized by increased microbial-derived lipopolysaccharide (LPS), potentially fueling the pathogenic effects of the estrogen metabolite 16α-hydroxyestrone (16α-OHE). These signals converge on macrophages, where co-exposure led to a hyperactivated state characterized by enhanced phagocytosis and ET-1 secretion. Tissue-level analyses confirmed immune cell infiltration and spatial association with elevated ET-1, providing evidence that these factors may contribute to the onset of sex-linked PAH. Taken together, these findings identify a previously unrecognized microbiome-estrogen-immune axis that amplifies BMPR2 dysfunction and provides a mechanistic basis for female-biased disease penetrance.

## I. Introduction

Sex differences in disease penetrance remain poorly explained by genetic variation alone. In pulmonary arterial hypertension (PAH), mutations in bone morphogenetic protein receptor type 2 (BMPR2) exhibit a marked female bias in disease development, despite worse outcomes in males, suggesting the involvement of sex-dependent modifiers beyond previously described canonical endocrine and chromosomal mechanisms.^1,2^ PAH is a cardiopulmonary disease characterized by vasoconstriction and vascular remodeling, which contribute to elevated right ventricular systolic pressure (RVSP) and right ventricular hypertrophy (RVH).^3^ PAH is progressive, and although current therapies improve symptoms, it remains with no cure. Patients often experience severe symptoms that result in right heart failure and, in many cases, death.^4^ PAH can arise from a variety of etiologies, with heritable PAH accounting for approximately 30% of all cases.^5^ Heritable PAH is caused by a number of mutations, the most prominent of which represent over 80% of heritable PAH cases, being mutations in the *Bmpr2* gene.^5^ Over 300 *Bmpr2* mutations have been identified in PAH patients,^6^ including the loss-of-function mutation R899X. Mutations in the *Bmpr2* gene have been shown to affect disease development and onset, while also eliciting a sex-linked predisposition previously described in the disease. ^7–9^ Specifically, females carrying the R899X mutation are four times more likely to develop PAH than their male counterparts.^8^ Despite the increased prevalence, women often have a higher survival rate. This is known as the estrogen paradox.^10,11^ The reasons for the sex disparity and the estrogen paradox remain under extensive discussion,^12^ with recent studies from our group demonstrating that endothelin-1 (ET-1), a potent vasoconstrictor, is markedly elevated in the lungs of humanized *Bmpr2^+/R899X^* mutant (MT) female animals.^7^

Elevated ET-1 expression has been associated with the development of PAH through its sustained vascular narrowing/vasoconstriction, contributing to elevated RVSP.^13,14^ Previous results have shown that macrophages, including Bone Marrow-Derived Macrophages (BMDM), play a significant role in the development of PAH. Specifically, Talati *et al.* (2010) reported that activated macrophages from BMPR2 MT produced more ET-1 than controls,^15,16^ but there was no further indication of sex-linked differences in PAH pathogenesis. Given that ET-1 promotes elevated vasoconstriction and a subsequent rise in RVSP, it is likely that increased ET-1 expression and secretion amplify the incidence in females with heritable PAH due to BMPR2-linked mutations. Investigating the role of ET-1 in this sex disparity, as well as understanding the source of this elevated expression, can be the start of a promising future for the development of a sex-specific treatment for those who suffer from PAH. Indeed, current therapies for PAH largely rely in part on ET-1 antagonists to mitigate its vasoconstrictor effects.^16^ Although these therapies improve patients’ quality of life, they are associated with adverse effects, such as peripheral edema and decreased hemoglobin.^17^ Additionally, currently clinically approved ET-1 antagonists, such as Bosentan and Ambrisentan, have been shown to be ineffective in mitigating chronic heart failure or reversing the progression of the disease.^17^ Thus, the search for viable alternative interventions remains a priority in the field of PAH research and medicine.

Although the mechanisms remain not entirely understood, gut microbiome dysbiosis has been linked to PAH.^18,19^ The homeostasis of the species that exist in the microbiome is essential for regulating the immune response. Similarly, Healthy lungs host a diverse community of microorganisms. ^20^ Like the gut microbiome, the lung microbiome can be altered by several factors, including infections, chronic lung diseases, and even mutations.^18,21^ As a result of lung microbiome dysbiosis, alterations in the presence of microbiome-derived metabolites can have implications for the disease.^22^ However, the link between the lung microbiome and the PAH sex disparity remains unclear. Within the lung microbiome, a collection of bacteria expressing ꞵ-Glucuronidase (ꞵ-Glu), called the estrobolome, exists. ꞵ-Glu is an enzyme that cleaves glucuronide molecules from conjugated estrogen metabolites, allowing for reabsorption of the estrogen metabolites instead of being excreted.^23,24^ Although ꞵ-Glu is important for physiological function due to its role in metabolic processes,^25^ dysbiosis of the microbiome can alter the function of the estrobolome, leading to increased ꞵ-Glu activity^24^ and the reabsorption of estrogen metabolites such as 16α-Hydroxyestrone (16α-OHE), a potent estrogen metabolite. 16α-OHE has been linked to an increase in the penetrance of BMPR2-associated PAH,^26^ meaning this metabolite increases the proportion of individuals with the genetic mutation who exhibit clinical characteristics of the described condition. It was demonstrated that not only was the predominant metabolism of estrogen to 16α-OHE associated with the penetrance of HPAH, but prolonged exposure contributed to the severity and penetrance of PH in *Bmpr2*-associated murine models.^27,28^ Although this harmful metabolite sheds light on the pathogenesis of sex-linked PAH, current therapeutic approaches remain challenging in part due to the lack of sex-linked specific molecular targets.^29,30^ In this context, elucidating the contribution of the lung estrobolome to PAH and its sex-linked disparity provides mechanistic insights and helps identify novel potential therapeutic targets, a central goal of this work.

## II. Methods and Materials

### 2.1. Animal Models

Male and female wildtype (WT) and *Bmpr2^+/R899X^* mutant (MT) mice were aged for 12 months and kept under normoxic conditions to allow spontaneous BMPR2 loss-of-function-associated progression of Pulmonary Hypertension (PH). A distinction is made in this study between PH and PAH, with PAH referring to the murine model; PAH is a term used for a clinical diagnosis that meets certain criteria in humans. After 12 months, mice were weighed and anesthetized with an intraperitoneal (IP) injection of Ketamine/Xylazine (K/X; 100 and 10 mg/kg, respectively). Lungs were perfused, then collected for metagenomic analysis; frozen in liquid nitrogen for subsequent western blot, protein measurement, and Enzyme-Linked Immunosorbent Assay; or fixed with 4% paraformaldehyde (PFA), sectioned for histological analysis.^7^ In all cases, strain-and age-matched mice were used as approved by the University of Illinois Chicago Institutional Animal Care and Use Committee.

### 2.2. Human Donors’ Lung Tissue

Deidentified human lung sections (formalin-fixed paraffin-embedded; tissue deemed unsuitable for transplant) were acquired from the Center for Heart Lung Innovation lung registry (Protocol Ethics No. H00-50110; University of British Columbia, Vancouver, BC, Canada).

### 2.3. Polymerase Chain Reaction (PCR)

*Bmpr2^+/R899X^* MT mice were genotyped *in-house* using an amplification refractory mutation system (ARMS)-modified PCR.^7^ Tails were collected in a sterile environment, and PCR was subsequently performed utilizing the REDExtract-N-AMP PCR kit protocol (Sigma, Cat #XnAT-100Rxn). Briefly, DNA was isolated from individually digested tails and used for genotyping along with the following primers: ARMS_mBMPR2_inFw: GAGGGAACGGCCATTAGAAGGTGGAT; ARMS_mBMPR2_outFw: AGAAGCCACAATGTTAATTCCCATGCTG; ARMS_mBMPR2_outrv: ATACTGCTGCCATCCAGGATATTTGTGG.

Samples were then placed in the PCR Thermal Cycler (Applied Biosystems, 2720 Thermal Cycler), products were resolved on a 1.5% agarose gel, and the gels were imaged using Li-Cor Odyssey CLx (Lincoln, NE) to determine the presence or absence of the MT fragment.

### 2.4. Lung Sample Preparation

Frozen perfused lung samples from male and female WT and *Bmpr2^+/R899X^*MT mice were homogenized at 4℃ with radioimmunoprecipitation assay (RIPA) buffer containing 1% protease inhibitor, and 0.1% phosphatase inhibitor cocktail 2 (#P5726), and 3 (#P004) purchased from Sigma Aldrich. Incubation was allowed for 20 minutes at 4℃, followed by 20 minutes of centrifugation at 12,000 *x*g (4℃). After centrifugation, the supernatant was collected, and protein concentration was measured using Bicinchoninic Acid (BCA) Assay as indicated below.

### 2.5. Bicinchoninic Acid Assay (BCA)

A standard concentration curve was prepared by serial dilution of Albumin (#23209; Thermo Scientific) to the following concentrations: 2, 1, 0.5, 0.25, 0.125, and 0.00 mg/mL (blank; milliQ water). Samples were prepared at a 10X dilution factor. Then, 5 µL of each sample was added *per* well of a 96-well plate; triplicate samples were used to account for error. 25 µL of reagent W (a mixture of reagent A and S) (#500-0113 and #500-0115, respectively; Bio-Rad), and 200 µL of reagent B (#500-0114; Bio-Rad), were added to each well, followed by 10-15 minutes of incubation. Intensity was measured using a microplate reader (Benchmark Scientific MR9600-T-SmartReader™) with a 495 nm filter, and samples were then interpolated from the standard curve using GraphPad Prism v9. Interpolated values were subtracted from the blank value.

### 2.6. Western Blot

After BCA determination, 20-30 µg of protein from the lung tissue lysate was diluted in Laemmli Sample Buffer (4x) with β-Mercaptoethanol, and samples were boiled at 95℃ for 10 minutes for Western Blot analysis. Before loading samples into a 1.5 mm SDS-PAGE gel, samples were centrifuged for 2.5 minutes at 16,200 *x*g. After running the samples through an 8%-12% gradient gel, protein separation was transferred onto a 0.2 µm nitrocellulose membrane. Samples were blocked with 5% milk or BSA (as specified on the datasheet) for 1 hour at room temperature. After unspecific blocking, the membrane was incubated with the primary antibody overnight at 4℃. The following day, the membrane was washed with TBS-Tween 1x (TBS-T: 2 x 10 min; 1 x 15 min), then incubated with a secondary HRP-conjugated antibody. After 1 hour of incubation, the membranes were washed again with TBS-T as indicated above. Membranes were then exposed to ECL reagent, and the chemiluminescence was scanned using Li-Cor Odyssey CLx. The expression level of the target protein was normalized using ꞵ-actin. Densitometric analysis was performed using ImageJ 1.54p.

### 2.7. Bone Marrow-Derived Macrophage (BMDM) Isolation

Intact femur and tibia isolated from male and female *Bmpr2^+/R899X^* MT mice were cleaned using sterile DMEM. Inside the biosafety cabinet, bone extremities were cut, and a 21-gauge needle was used to push 5 mL of DMEM through the bone marrow. Cells were collected in a 50 mL Falcon tube, counted, and 10^6^ cells *per* well were plated onto a 6-well plate. BMDM underwent differentiation for 7 days through exposure to 10% L929 supernatant. Cell morphology was observed, and medium was replaced approximately every 48 hours.^31^ On day 7, the cells were treated with either vehicle control, 100 ng/mL Lipopolysaccharide (LPS), 100 nM 16α-OHE, or a cotreatment containing both 100 ng/mL LPS and 100nM 16α-OHE. After 24 hours, the cell supernatant was collected, and cell lysates were prepared using RIPA buffer plus 1% protease inhibitor and 0.1% phosphatase inhibitor as described above. Alternatively, cells were also used to perform a phagocytosis assay as indicated below.

### 2.8. Phagocytosis Assay

Differentiated BMDM were treated with vehicle control, 100 ng/mL LPS, 100 nM 16α-OHE, or 100 ng/mL LPS + 100 nM 16α-OHE for 24 hours, when cells were exposed to 10^7^ particles/well of Fluorescein isothiocyanate (FITC)-labeled heat-inactivated *Escherichia coli* (E13231; Invitrogen) for 1 hour.^31^ Then, 10^5^ cells/well were transferred to a black 96-well plate with a transparent bottom, and fluorometric measurements were immediately performed using BioTek Synergy H4 Hybrid Multi-Mode Microplate Reader (Settings: 485/20, 528/20, Gain 100). Alternatively, cells were used for immunocytochemistry.

### 2.9. Immunocytochemistry (ICC)

10^2^ BMDM were transferred to Lab-Tek Chamber Slide System and quenched with 1:2 trypan blue in PBS for 1 minute. The cells were then fixed using 4% paraformaldehyde (PFA). The cells were permeabilized and blocked with 1% BSA for 30 minutes. After blocking, Primary intubation was performed with ET-1. The slide was washed 3x with 0.025% Triton X-100 before incubation with the secondary antibody Alexa 555. Vectashield containing DAPI was applied to the slide before a coverslip was placed. Images were taken using an LSM880 confocal microscope (Carl Zeiss Microimaging, Inc.).

### 3.0. Immunohistochemistry (IHC) and histological analysis

PFA-fixed, paraffin-embedded lung tissue sections of male and female *Bmpr2^+/R899X^* MT mice were used to observe and quantify protein expression in specific regions of the lungs. Deparaffinization was performed on the slides before immunostaining through serial exposure to Xylene and Ethanol. Antigen retrieval was performed in sodium citrate buffer at high temperature and pressure for 10-15 minutes. Slides were washed with 0.025% Triton buffer (2x 5 min) and then blocked with 5% BSA for 1 hour at room temperature. After blocking, the slides were incubated with the primary antibody overnight at 4℃. The following day, the slides were washed again with a Triton buffer (2x 5 min). The slides were then incubated with Alexa fluorescence-labeled secondary antibody for 1 hour at room temperature. The slides were then washed again with a Triton buffer (2x 5 min). Vectashield containing DAPI was used to stain nuclei. Micrographs were obtained using an LSM880 confocal microscope (Carl Zeiss Microimaging, Inc.). Additionally, Hucker-Twort staining was performed to distinguish Gram-positive (green/blue) and Gram-negative (red/pink) bacteria in lung tissue sections using a commercially available kit (Gram, Hucker-Twort Stain Kit; Newcomer Supply, Inc., Madison, WI, USA) according to the manufacturer’s instructions. Slides were scanned using an Aperio brightfield automated microscope slide scanner (Leica Aperio AT2) and micrographs analyzed using ImageScope software 12.4.6 (Leica Biosystems).

### 3.1. Enzyme-Linked Immunosorbent Assay (ELISA)

Frozen perfused lung samples were homogenized in RIPA solution as previously described and used to quantify levels of LPS (EU3126; FineTest), Lipoteichoic acid (LTA) (ELK8790; ELK Biotechnology), Peptidoglycan (MBS755183), and 16α-OHE (MBS263184) from MyBioSource, according to the manufacturer’s protocol.

### 3.2. Flow Cytometry Analysis

Freshly isolated lung samples from male and female WT and *Bmpr2^+/R899X^*MT mice were washed with cold PBS before being finely minced with a razor blade in 500 µL DMEM. 8.5 mL more of DMEM was added to the mixture, followed by 1mL of collagenase type I (20 mg/mL), bringing the final volume to 10 mL. Samples were incubated for 30 minutes at 37℃ with agitation. After incubation, samples were passed through a 0.45 µm filter. The samples were then spun down at 350 ×g for 5 minutes before resuspension in Hank’s balanced salt solution (HBSS). For flow cytometry analysis, 10^5^ cells were transferred to a well at 96 well-plate, spun down at 350 *x*g for 5 minutes, resuspended in 20 µL of staining buffer, incubated with 5 uL Fc receptor block solution (5 minutes; 4°C) and stained with 0.25 µL of FITC anti-F4/80, 1.25 µL of PE-Cy7 anti-CD45 and 0.3 µL of APC anti-CD31 for 20 min at 4°C. After surface stain, cells were washed twice with 150 µL of staining buffer and permeabilized for 10 min at 37 °C using Fix Buffer I. Cells were then washed twice and resuspended in 25 µL of staining buffer and incubated with 0.5 µL of anti-ET-1 antibody for 20 min at 4°C. After incubation, cells were washed twice, resuspended in 25 µL of staining buffer, and stained with 0.5 µL of PE anti-mouse. After intracellular staining, cells were washed twice and resuspended in 150 µL of staining buffer for flow cytometry analysis on a Cytek Northern Light cytometer (Cytek^®^ Biosciences). Unstained cells and fluorescence minus one (FMO) controls for each antibody were used. Dot plots were generated using SpectroFlow Jnk or FlowJo™ Software.

### 3.3. Lung Metagenomic Analysis

Lung tissue was collected from WT and *Bmpr2^+/R899X^* MT mice of both sexes for whole metagenomic analysis. The samples were placed in bar-coded collection tubes containing DNA stabilization buffers to preserve DNA integrity. Samples underwent whole-genome sequencing (WGS) by TransnetYX. After sequencing, low-quality or mislabeled records were removed before analysis using a combination of automated and manual approaches. Quality control also involves screening out host rodents and human genomes before raw data alignment with the most recent One Codex Microbiome Database. Aligned data were used to determine the microbiota richness and evenness in the samples (*i.e*., α-diversity) and the similarity or dissimilarity between communities (*i.e.,* β-diversity). Metagenomic classification readcounts_with_Children was used. Data were only analyzed when the readcount exceeded 30K reads based on our negative PBS control. Then, samples were examined at the Phylum, Genera, and species levels, and the most frequent estrobolome genera and species were plotted and statistically compared using GraphPad Prism v9, whereas the relationship between sex, genotype, and genera was analyzed using RStudio. The remaining reads were listed as “Others” and often represent < 30% of the readcounts. Raw data analysis was conducted using One Codex analysis software (https://onecodex.com/, latest access date: 18 March 2024). Adherence to the data-sharing policies will be ensured by depositing FASTQ files in the NCBI Sequence Read Archive and by depositing codes on GitHub (github.com/OliveiraSD/VI Lab_2024), which will be publicly available upon peer-reviewed publication.^18^

### 3.4. Liquid chromatography-mass spectrometry (LC-MS) for targeted metabolomics

Short-chain fatty acids (SCFAs) were isolated from plasma and frozen pulverized lung lysates. Briefly, 5 μL of the calibrator and sample were injected into an AB SCIEX 5500 QTRAP coupled with an Agilent 1290 UPLC system (samples eluted by Agilent Poroshell column 120 EC-C18 2.7 μm, 2.1 x 100 mm with a flow rate of 450 μl/min; column compartment at 40 °C). LC elution was 99% mobile phase A (0.1% FA in H2O) for 1 min, followed by a linear gradient increase of mobile phase B (0.1% FA in ACN) from 1% to 10% in 1 min, then from 10% B to 65% B in 6 min and from 65% B to 90% B in 0.1 min. The column was washed with 90% B (3 min), then re-equilibrated to the initial condition (99% A; 3 min). The autosampler was maintained at 4 °C. MS data were acquired by MRM scan in negative mode. The ESI spray voltage and source temperature were set to −4.5 kV and 450 °C, respectively. Negatively charged analytes and standards were detected by monitoring their transition to signature product ions.

### 3.5. Reagents and Antibodies

Rabbit polyclonal ET-1 (#12-191-1-AP) was obtained from ProteinTech. Unconjugated monoclonal Rabbit CD45 antibody was obtained from Cell Signaling Technology (#702575). Rabbit GUSB antibody was obtained from Protein Tech (16332-1-AP). eNOS was obtained from Santa Cruz Biotechnology (sc-376751). F4/80 was obtained from Abcam (ab300421). Polyclonal Goat anti-Rabbit Alexa 555 secondary antibody was obtained from Invitrogen (#A21429). Polyclonal Donkey anti-Rabbit Alexa 555 secondary antibody was obtained from Invitrogen (A-31572). Polyclonal Donkey anti-Goat Alexa 488 was obtained from Invitrogen (A11055). Monoclonal Mouse-HRP was obtained from Bio-Rad (#**1706516**). Anti-Rabbit IgG (H+L) antibody peroxidase-labeled was obtained from SeraCare (5220-0283). Mouse Anti-NHP CD44 (561294), Fix Buffer I (Cat# 557870), and PE-Cy7 anti-CD45 (Cat# 561294) from BD Bioscience; APC Anti-mouse CD31 Antibody (Cat# 102510), staining buffer (Cat# 420201), Fc receptor block solution (Cat# 422302) from BioLegend; F4/80 Monoclonal Antibody (BM8), FITC (Invitrogen, 11-4801-82), and PE Polyclonal anti-mouse IgG (Cat# 554047) from BD **(Table 1).**

### 3.6. Statistical Analysis and Rigor

Data was analyzed utilizing GraphPad Prism v10. Normality was determined through the Shapiro-Wilk test. Data was plotted as mean ± Standard Error Mean (SEM). Normally distributed data were analyzed using a two-way ANOVA with a *post hoc* test (Bonferroni) or a Parametric (Student’s t-test) analysis. Data that were not normally distributed were analyzed using a non-parametric test (Mann-Whitney test) to determine differences between two or more groups. *P* < 0.05 was considered significant. IHC results for lung CD45^+^ cells and β-glu expression were analyzed in a double-blinded manner. Specifically, one individual performed the immunostaining, and another collected confocal micrographs of the pulmonary microvasculature, without knowing the animal’s sex. Finally, a third individual recorded the perivascular cells or fluorescence intensity in each sample, unaware of the animal’s sex. Microvessels were outlined using the spline tool either along the endothelial cell lining or around the perivascular space, and the mean fluorescence intensity of the desired channel was measured using Zeiss Zen 3.7 Software.

## III. Results

### 3.1. Perivascular CD45^+^ cells may contribute to lung ET-1 rise in female *Bmpr2^+/R899X^*

Recently, we reported that humanized *Bmpr2^+/R899X^* female MT animals displayed significantly high lung ET-1 expression and RVSP at 12 months of age, despite mild vascular pulmonary vascular remodeling.^7^ When investigating the source of elevated lung ET-1 expression, data indicated that carrying the R899X mutation resulted in increased colocalization of CD45 and ET-1 expression in the microvasculature of female MT. However, this pattern was not observed in their male counterparts, indicating a putative sex-specific association between immune cell infiltration and increased ET-1 expression in the lungs of the *Bmpr2^+/R899X^* animal model **(Fig. 1A)**. Consistent with this observation, increased ET-1 expression in whole lung tissue from *Bmpr2^+/R899X^* MT females^7^ is not observed between 12 month-old males and females expressing normal level of BMPR2 protein (WT) **(Fig. 1B)**. This data indicates that carrying the human loss-of-function *Bmpr2^+/R899X^*mutation drives a female-biased increase in pulmonary ET-1 levels, known to be essential in development of PAH. To assess the extent of perivascular immune cell accumulation in MT males and females, the pan-leucocyte marker CD45 was used, and CD45^+^ cells were quantified from lung IHC micrographs, revealing that MT females’ microvessels contained significantly more CD45^+^ cells than those of male MT **(Fig. 1C-D)**. These findings indicate an association between perivascular immune cell presence and ET-1 expression in the lungs of female MT animals.

**Figure 1:**
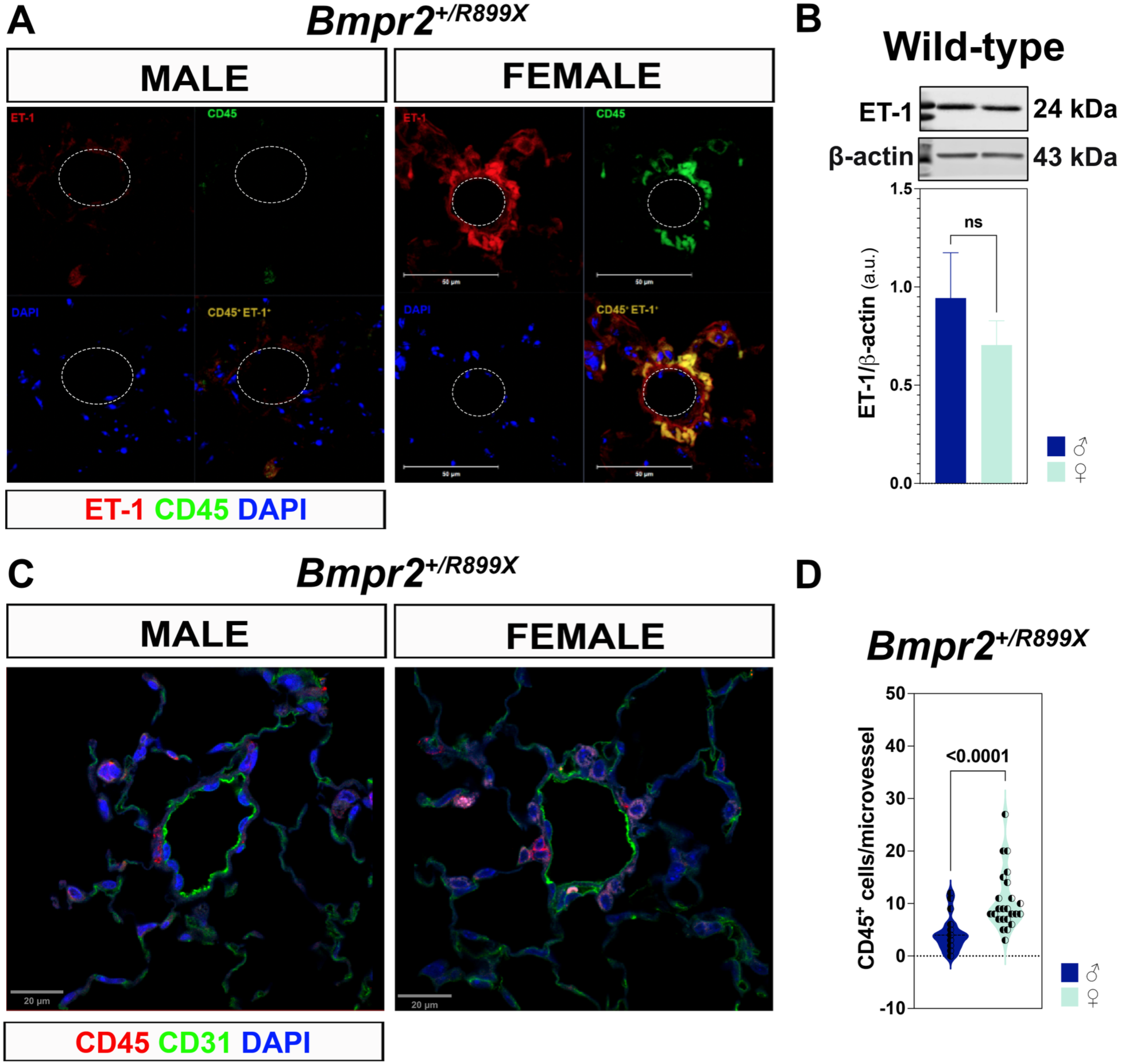
Increased perivascular CD45+ cell accumulation colocalizes with increased endothelin-1 expression in pulmonary microvessels of female humanized *Bmpr2^+/R899X^*mice. 12-month-old *Bmpr2^+/R899X^* mutant male (left) and female (right) lung tissue was evaluated using immunohistochemistry (IHC), showing expression of endothelin-1 (ET-1) (red), CD45 (green) in microvessels (dashed white lines; < 100 μm). DAPI (blue) was used to stain cell nuclei. Staining suggests colocalization of the pan-leukocyte marker CD45 and ET-1 expression (orange) in MT females **(A)**. Lung tissue was used to analyze ET-1 expression in male (♂; dark blue) and female (♀; cyan) wild-type mice by Western blot. ꞵ-actin was used as a loading control **(B)**. IHC analysis showing expression of CD45 (red), CD31 (green), and DAPI (blue) in the pulmonary microvessels of male (left) and female (right) *Bmpr2^+/R899X^* mutant mice. Micrographs were used to quantify the number of CD45+ cells in pulmonary microvessels (replicates = 6) **(C, D)**. All microvessels imaged had a diameter of no greater than 100 µm. IHC images were obtained by confocal microscopy, and quantification was performed by a blinded investigator for animal sex. The Shapiro-Wilk test was used to determine whether the data were normally distributed. Normally distributed data were analyzed using two-way ANOVA followed by Bonferroni *post hoc* test (n = 3-4 animals per group). Data is represented as mean ± SEM (ns = non-significant).

### 3.2. Sex-linked lung macrophage abundance and ET-1 expression in *Bmpr2^+/R899X^* mice

Consistent with findings in Fig. 1, IHC imaging revealed that the CD45^+^ leukocytes around the microvasculature of MT female animals are mainly F4/80^+^ cells, indicating that macrophages likely contribute to the prevalent perivascular immune cell population in early PAH onset **(Fig. 2A).** To further characterize this population and also, taking into consideration the largely known contribution of vascular endothelial cells to ET-1 level, flow cytometry was performed to characterize both CD45^+^F4/80^+^ and CD45^-^CD31^+^ lung cells **(Fig. Suppl. 1A)**. Using a freshly isolated single-cell suspension of lung cells, flow cytometry analysis revealed the presence of both F4/80^high^ and F4/80^low^ subsets, consistent with tissue-resident macrophages and recruited monocyte-derived macrophages, respectively.^31,32^ Additionally, F4/80^+^ lung cells from female *Bmpr2^+/R899X^* MT had a significantly greater proportion of the F4/80^low^ population than male MT animals; however, there was no observable difference in F4/80^high^ expression among groups **(Fig. 2B)**. Flow cytometry analysis of intracellular ET-1 expression revealed no significant sex-dependent differences in the overall proportion of ET-1-positive lung endothelial cells or macrophages **(Fig. 2C-F)**. Given that ET-1 is a secreted peptide, intracellular detection by flow cytometry provides a snapshot of cell-associated ET-1 and should not be interpreted as a direct measure of extracellular ET-1 abundance or release. Together, these findings indicate that early disease in females carrying the *Bmpr2^+/R899X^* mutation may be characterized by a shift toward a more inflammatory, monocyte-derived macrophage phenotype, accompanied by enhanced ET-1-mediated signaling, while endothelial ET-1 expression remains unchanged. These observations support a putative model in which macrophage heterogeneity and endothelial-immune crosstalk may contribute to the sex-biased immune-vascular interaction underlying early PAH development.

**Figure 2:**
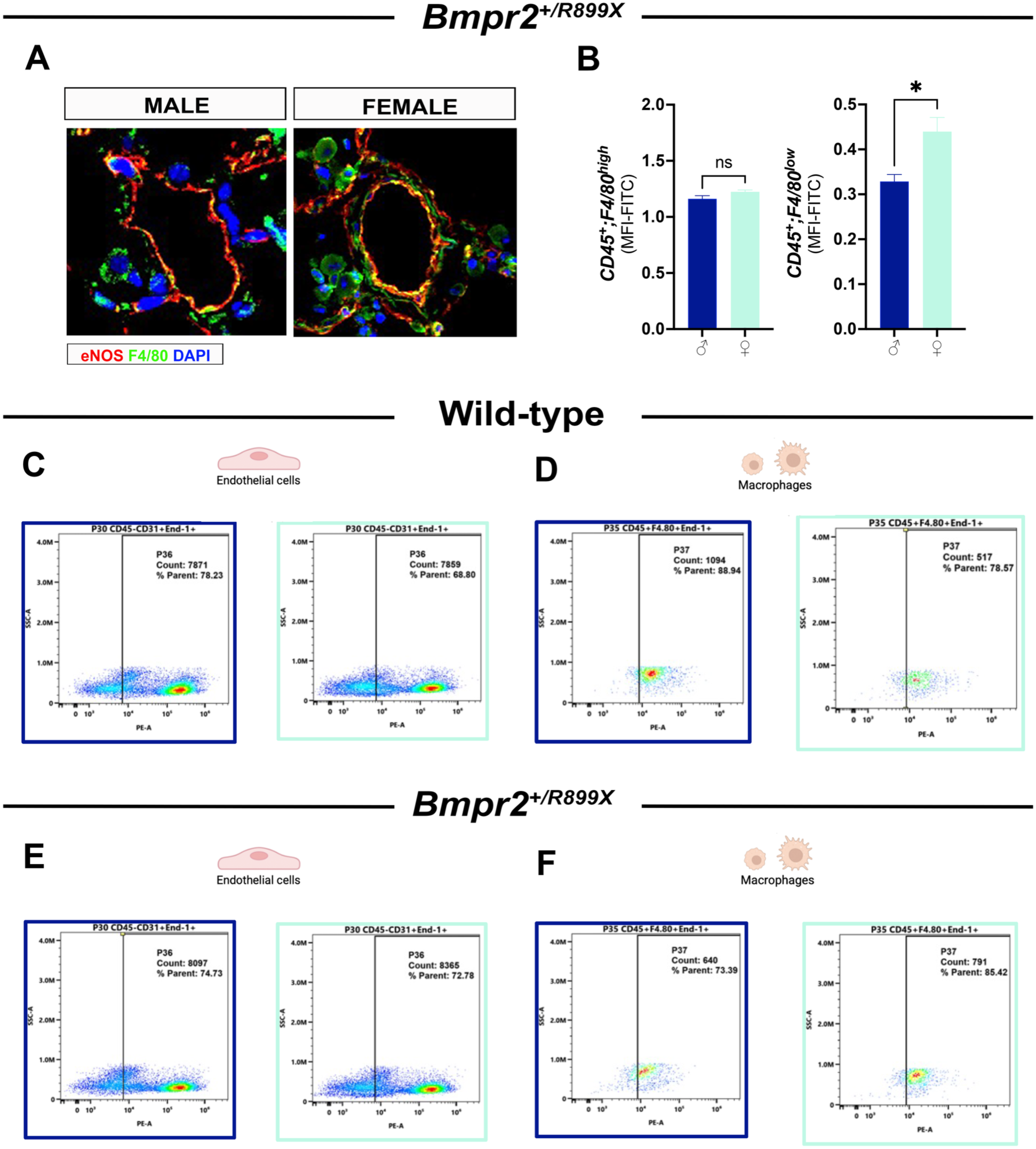
Sex-linked differences in pulmonary macrophage abundance and endothelin-1 expression in humanized *Bmpr2^⁺/R899X^* mice. Immunohistochemistry (IHC) was performed on lung sections from 12-month-old male and female *Bmpr2^+/R899X^* mutant (MT) mice, with pulmonary microvessels stained for eNOS (red), F4/80 (green), and DAPI (blue) **(A)**. Representative images show increased perivascular localization of F4/80⁺ macrophages in females compared to males. Quantification by flow cytometry demonstrates the mean fluorescence intensity (MFI) of CD45⁺F4/80^high^ and CD45⁺F4/80^low^ populations in male (dark blue) and female (cyan) MT mice, normalized to FITC-F4/80 intensity **(B)**. Representative flow cytometry plots from lung single-cell suspensions are shown for wild-type (WT) and *Bmpr2^+/R899X^* mice. Endothelial cells (CD45⁻CD31⁺) and macrophages (CD45⁺F4/80⁺) were gated, and endothelin-1-positive cells were quantified in WT **(C-D)** and MT **(E-F)** animals. Data are representative of n = 5-6 animals per group. Statistical significance was assessed using Student’s t-test, with *p < 0.05 and ns indicating not significant.

### 3.3. ET-1 increases in *Bmpr2^+/R899X^* Female BMDM as a Response to Microbial Molecules

To assess the contribution of sex to non-resident macrophage function, specifically their response to potential microbial threats, BMDMs isolated from male and female animals were treated with 100 ng/mL LPS or vehicle control. Results demonstrated not only that BMDM from *Bmpr2^+/R899X^* MT females expressed higher ET-1 levels upon LPS stimulation, but also that they secreted significantly more vasoactive peptide than their male counterparts **(Fig. 3A, B, C)**. Predisposition to LPS-induced ET-1 secretion by macrophages from *Bmpr2^+/R899X^*MT females prompted an investigation into the presence or absence of Gram-negative bacteria in their lungs. IHC analysis confirmed that both Gram-positive and Gram-negative bacteria are present in WT and MT lung tissue **(Fig. 3D)**. Interestingly, Gram-negative bacteria were observed mainly within the pulmonary microcirculation, whereas Gram-positive bacteria were more widespread in the *Bmpr2^+/R899X^* MT lungs **(Fig. 3E)**. Corroborating with histological analysis, ELISA indicated that LPS levels were significantly higher in the lungs of *Bmpr2^+/R899X^* MT females than in males, signifying the presence of Gram-negative bacteria. On the other hand, evaluation of peptidoglycan and LTA, major components of gram-positive bacteria,^33^ revealed no significant difference between MT males and females **(Fig. 3F-H)**. Data suggest that the immune response of monocyte-derived macrophages in *Bmpr2^+/R899X^* MT females is likely elicited by continued low-level exposure to Gram-negative bacteria in the lungs and is also likely responsible for inducing localized ET-1 secretion in the microcirculation.

**Figure 3:**
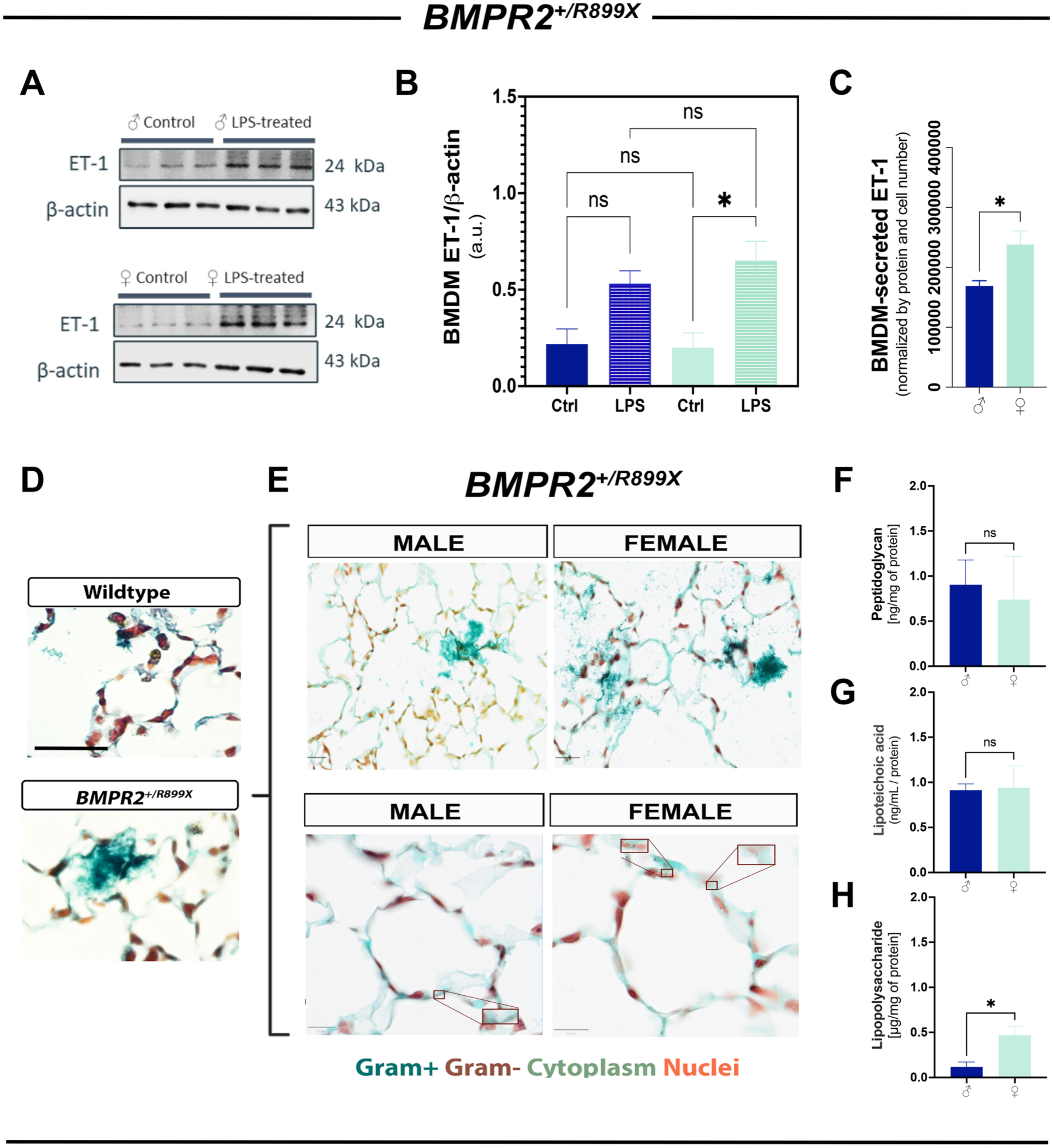
Activation of *Bmpr2^+/R899X^*mutant female bone marrow-derived macrophages in response to antigenic molecules. *In vitro* analysis of Bone Marrow-Derived Macrophages (BMDM) isolated from 12-month-old male and female *Bmpr2^+/R899X^* mutant (MT) animals was performed, with cells treated with either vehicle control or 100 ng/mL Lipopolysaccharide (LPS) for 24 hours. Cell lysates were analyzed by western blot to show endothelin-1 (ET-1) expression in male (dark blue) and female (cyan) *Bmpr2^+/R899X^* MT animals. ꞵ-actin was used as a loading control **(A)**. Cell Supernatant was used to detect secreted ET-1 expression in control male and female *Bmpr2^+/R899X^*MT by dot blot analysis. Data were normalized by protein concentration and cell number **(B)**. Hucker-Twort staining was performed to evaluate Gram-positive (green) and Gram-negative bacteria (red) in the lungs of wild-type and *Bmpr2^+/R899X^* MT **(D, E).** The presence of Gram-negative bacteria was also assessed by measuring LPS levels in lung lysates. Data was normalized with protein concentration **(F)**. Quantification of Gram-positive bacteria was analyzed using two methods: ELISA for Peptidoglycan and Lipoteichoic acid (LTA). Both methods were normalized with protein concentration of the cell lysates **(G, H)**. Normality was determined through the Shapiro-Wilk test. Normally distributed data were analyzed using Student’s t-test or two-way ANOVA followed by Bonferroni post hoc test. Data is represented as mean ± SEM (n=3-4 per group). *p<0.05, ns, non-significant.

### 3.4. *Bmpr2^+/R899X^* Mutation Contributes to Sex-linked Lung Microbiome Dysbiosis

Differential observations of bacteria within the lung tissue prompted further investigation of the lung microbiome. Then, after metagenomic analysis of perfused lung tissue from WT animals and *Bmpr2^+/R899X^* model, we assessed α-diversity using Shannon, Simpson, and Chao1 indices, and ꞵ-diversity using unweighted UniFrac. Among WT animals, there were no significant differences between males and females in each of the α-diversity indices **(Fig. 4A-C).** Additionally, unweighted unifrac revealed overlapping and comparable community composition **(Fig. 3D)**. Conversely, when assessing MT animals, females displayed elevated species richness in the Chao1 index **(Fig. 4E-G)**. Unweighted unifrac also revealed distinct separation in community structure between males and females **(Fig. 4H)**, suggesting that the *Bmpr2^+/R899X^* mutation results in a sex-specific alteration of the lung microbiome and providing compelling evidence that MT females are undergoing lung microbiome dysbiosis. Moreover, when analyzing the four most abundant phyla, the % frequencies of Bacteroidetes, Firmicutes, Ascomycota, and Proteobacteria did not differ significantly between WT males and females **(Fig. 4I-M)**. However, analysis among *Bmpr2^+/R899X^* MT males and females revealed a minor yet significant difference in the % of Ascomycota **(Fig. 4N-R)**. Given that microbiome dynamics significantly regulate systemic and local metabolites with inflammatory properties, including short-chain fatty acids (SCFAs), targeted metabolomics was performed using liquid chromatography-mass spectrometry (LC-MS). LC-MS analysis of WT and MT-derived plasma and lung samples revealed that all four major SCFAs analytes (*i.e.,* acetate, propionate, butyrate, and valerate) were significantly higher in the plasma compared to lung tissue, but not statistically different between WT and MT. Acetate was most abundant in MT than in WT, with propionate, butyrate, and valerate exhibiting opposing levels. Additionally, male and female WT mice exhibited no difference in plasma SCFAs, whereas all metabolites were increased in the WT female lungs compared to WT males. On the other hand, sex-associated differences in the plasma or lung levels of all SCFAs were absent in *Bmpr2^+/R899X^* MT mice (**Fig. Suppl. 2A-E)**. This data continues to provide convincing evidence of sex-specific dysbiosis of the lung microbiome in individuals carrying the *Bmpr2^+/R899X^* mutation. Human studies would be critical to further evaluate the translational potential of these findings.

**Figure 4:**
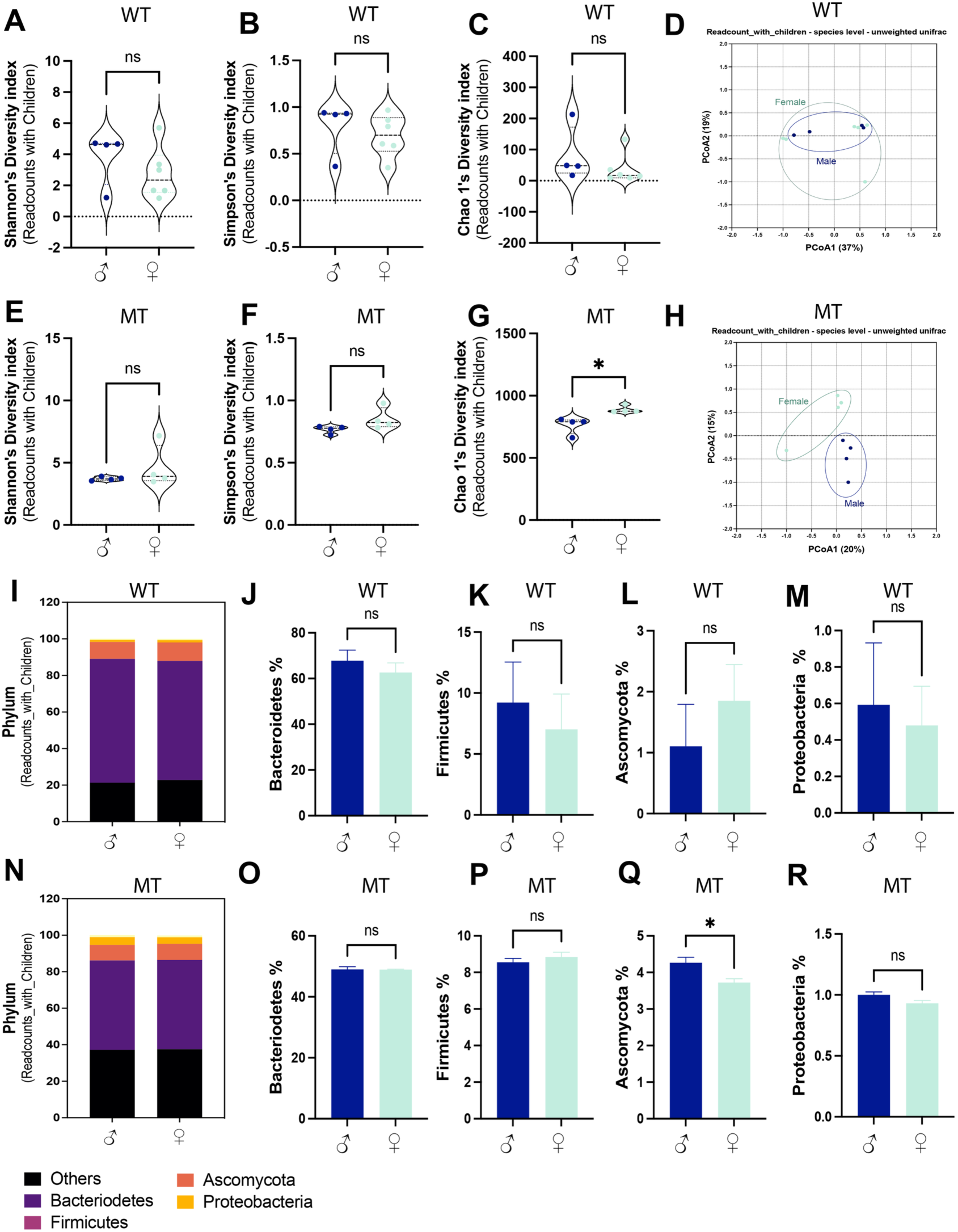
Lung metagenomic analysis of 12-month-old male and female Wildtype and *BMPR2^+/R899X^*humanized animals. Whole genome sequencing was performed in lung tissue from male (dark blue) and female (cyan) 12-month-old male and female Wildtype (WT) and *Bmpr2^+/R899X^*mutant (MT) animals. α-diversity analysis was performed using Shannon, Simpson, and Chao1’s diversity indices **(A-D, E-G).** ꞵ-diversity was analyzed using unweighted unifrac as shown in the Principal Component Analysis (PCA) plot **(D, H).** Readcount % of the top four most frequent Phyla in WT and *Bmpr2^+/R899X^* MT. Bacteroidetes (purple), Ascomycota (orange), Proteobacteria (yellow), Firmicutes (violet), Other (black) **(I, N).** Additional graphs showing comparison between males (dark blue) and females (cyan) in each of the top four Phyla **(J,-M, O-R).** Readcounts with children were used to compare the lung microbiome among groups. The Shapiro-Wilk test was used to determine normality. Parametric (Student’s t-test) or nonparametric (Mann-Whitney test) tests were used based on the results of the normality test. Data were analyzed using RStudio and One Codex Cloud Platform (n=8-10 animals per group). *p<0.05, ns, non-significant.

### 3.5. Lung Estrobolome May Contribute to Accumulation of Cytotoxic 16α-OHE Metabolite

Sex-linked changes in the lung microbiome guided an investigation into how microbes would functionally contribute to the *“sex paradox”* observed in PAH. It is known that the microbiome harbors bacteria that play a key role in hormonal regulation by promoting the reabsorption of estrogen and its metabolites via β-glu-mediated mechanism. This bacterial group, the estrobolome, was identified in lung tissue from both WT and MT animals, but in the MT group, increased *Lactobacillus* and *Eubacterium* contrasted with reduced *Clostridium*, *Faecalibaculum, Alistipes*, and *Prevotella* **(Fig. 5A)**. Chord diagram analysis of the sex-linked estrobolome genera revealed a predominance of *Lactobacillus* and *Eubacterium* in *Bmpr2^+/R899X^* MT females **(Fig. 5B)**. Previously reported as toxic in PAH, the estrogen metabolite 16α-OHE was also found to be significantly elevated in the lung tissue of MT females (**Fig. 5C)**. ꞵ-Glu expression was also analyzed in order to evaluate the putative functional contribution of the lung estrobolome in the accumulation of 16α-OHE. Although ELISA analysis of whole lung lysate did not reach a statistically significant difference between sexes **(Fig. 5D)**, IHC analysis of lung ꞵ-Glu intensity revealed that female MT have significantly greater ꞵ-Glu localized in the inner endothelial lining of the microvasculature with the presence of ꞵ-Glu^+^ perivascular cells **(Fig. 5E-G)**. Similarly, initial confocal imaging of the remodeled pulmonary microvessels in one female and one male patient corroborates our findings in the humanized PH animal model **(Fig. 5H)**. Finally, analysis of ꞵ-Glu in human endothelial cells, macrophages, and alveolar cells from publicly available datasets in Human Protein Atlas^34^ (proteinatlas.org) revealed no significant sex-based difference between healthy males and females among these cell types **(Fig. Suppl. 3)**. Altogether, this data suggests that estrobolome may contribute to elevated ꞵ-Glu presence in the endothelial lining of blood vessels in females carrying the *Bmpr2* mutation or with deficiency in lung BMPR2 expression, which corroborates with increased unconjugated 16α-OHE level observed in the female MT lungs.

**Figure 5:**
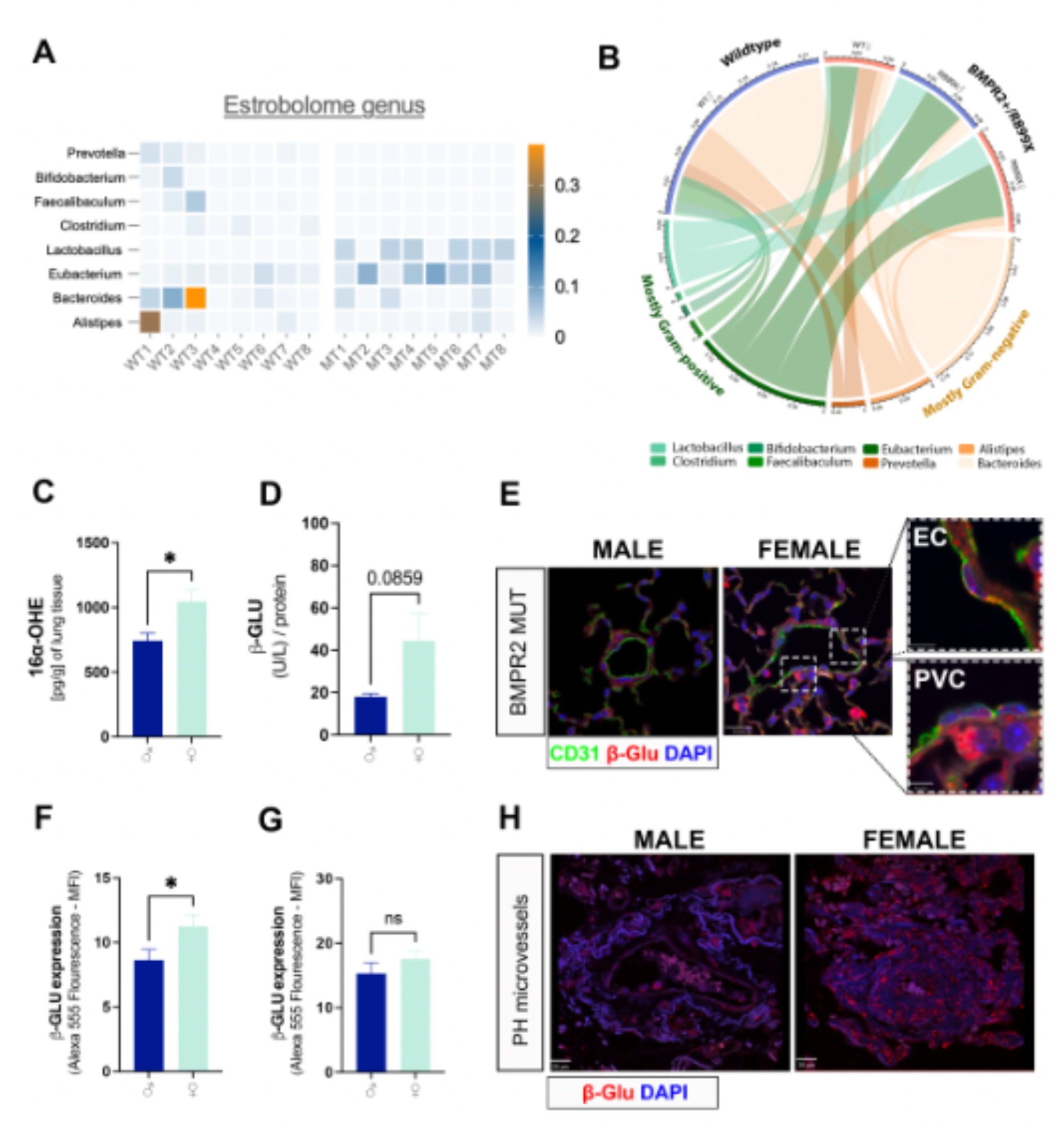
Lung Estrobolome and sex-linked accumulation of cytotoxic estrogen 16α-OHE metabolite. The panel shows the estrobolome microbial genera predominance in lung tissue of 12-month-old male and female Wildtype (WT) and *Bmpr2^+/R899X^* mutant (MT) animals **(A).** The estrobolome associations identified in the chord diagram are shown in green (gram-positive) and orange (gram-negative), along with the genera comprising the populations **(B)**. ELISA analysis showing concentration of 16α-Hydroxyestrone (16α-OHE) and ꞵ-Glucuronidase (ꞵ-Glu) expression in the lungs of male (dark blue) and female (cyan) *Bmpr2^+/R899X^* MT mice **(C-D)**. Representative IHC images of the pulmonary microvasculature, stained for CD31 (green), β-Glu (red), and DAPI (blue) in *Bmpr2^+/R899X^* MT males and females **(E)**. Additionally, IHC analysis of mean intensity value of Alexa 555, representing ꞵ-Glu expression, bordered around the endothelium **(F)** as well as bordered around the perivascular surround of the microvasculature (n=3) (replicates=10) **(G)**. IHC analysis of Alexa 555 mean intensity, representing ꞵ-Glu expression, was performed on lung sections from one male and one female human PH patient donor **(H)**. The Shapiro-Wilk test was used to determine whether the data were normally distributed. Parametric (Student’s t-test) or nonparametric (Mann-Whitney test) tests were used based on the results of the normality test. (n=3-4 animals per group) *p<0.05, ns, non-significative. EC = endothelial cell; PVC = perivascular cell.

### 3.6. 16α-OHE Increased Phagocytosis and ET-1 Production in BMDM from Female MT

Increased estrobolome-linked ꞵ-Glu and perivascular F4/80^low^ macrophage accumulation in the pulmonary circulation, along with elevated LPS and 16α-OHE levels in females carrying the *Bmpr2* mutation, raised questions about the role these molecules play in BMDM activation in response to bacterial exposure and subsequent ET-1 production. Immunostaining of ET-1 in BMDM exposed to FITC-tagged *E. coli* suggested that, individually, cells that phagocytosed more *E. coli* also expressed less intracellular ET-1 level **(Fig. 6A)**. Quantitative analysis of the overall phagocytosis activity revealed significantly greater spontaneous BMDM activity in female MT-derived macrophages, but this difference is lost when cells were exposed to 16α-OHE for 24 hr. Interestingly, when BMDM from female MT were co-incubated with both LPS and 16α-OHE, phagocytosis increased substantially at the same time ET-1 secretion was also elevated, compared to male-derived cells **(Fig. 6B, C)**. These findings suggest that, *in vivo*, accumulated lung 16α-OHE may synergize with Gram-negative bacteria, contributing to increased female RVSP^7^ by leading recruited macrophages to oversecrete ET-1 on the microcirculation, which in turn may potentiate dedifferentiation of vascular cells into a myofibroblast-like contractile and hyperproliferative phenotype^7^, a process that may occur through Endothelial-To-Mesenchymal transition (EndoMT) (Wermuth), contributing to sex-linked disease onset **(Fig. 7)**.

**Figure 6:**
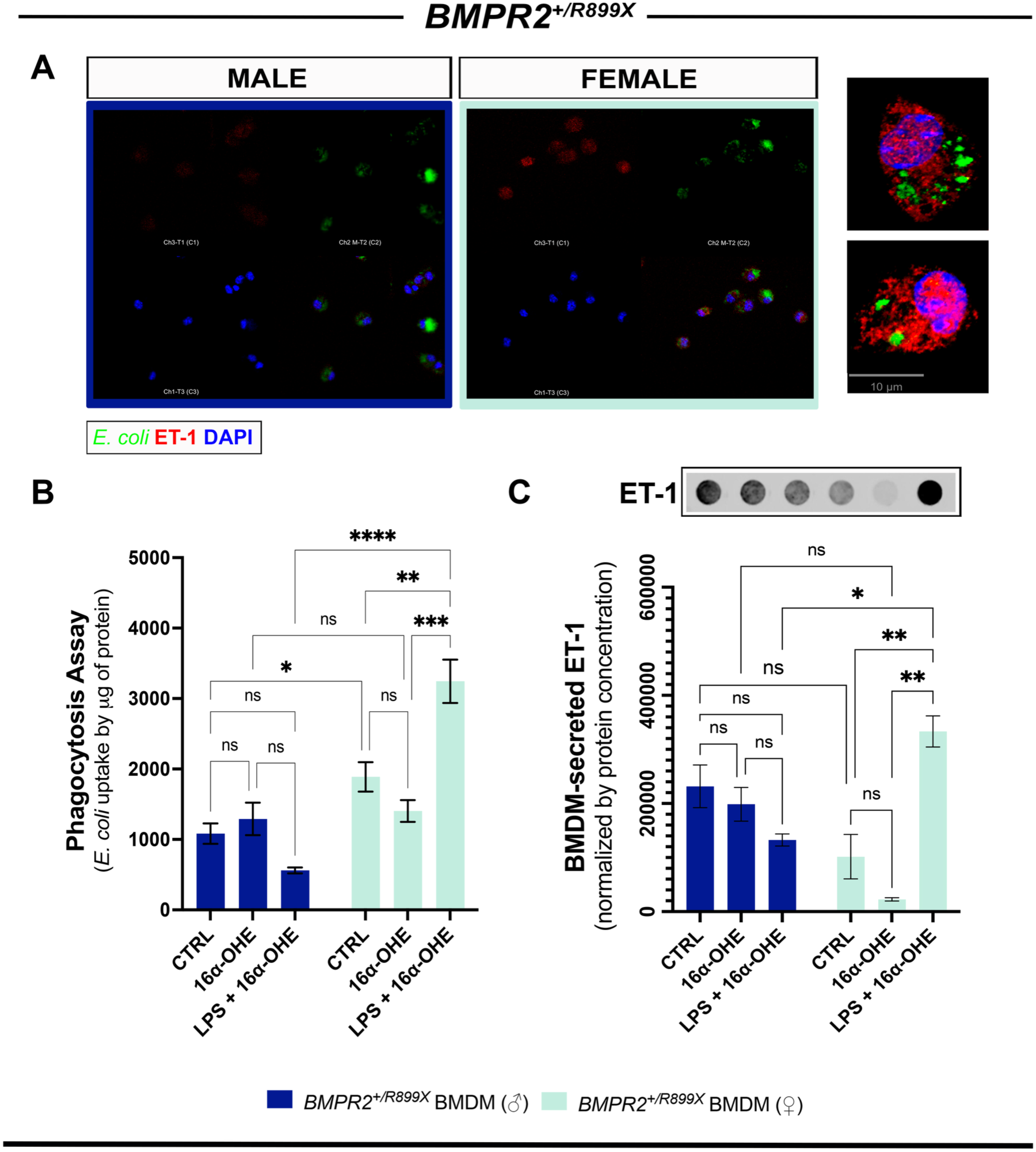
LPS and 16α-OHE synergistically increased macrophage phagocytosis and ET-1 production in Female *Bmpr2^+/R899X^* animals. Immunocytochemistry showing the phagocytosis of FITC-tagged *E. coli* (green), Endothelin-1 (ET-1), and DAPI (blue) in the Bone Marrow Derived Macrophages (BMDM) of male (left) and female (right) *Bmpr2^+/R899X^* mutant (MT) mice **(A)**. *Inset:* A higher magnification image shows BMDM with less ET-1 and more *E. coli* (top) and more ET-1 and less *E. coli* (bottom). Phagocytosis Assay showing the intake of fluorescent tagged *E. coli* in BMDM from MT male (dark blue) and female (cyan) mice after treated for 24 hours with a vehicle control, 100 ng/mL of Lipopolysaccharide (LPS), 100 nM of 16α-Hydroxyestrone (16α-OHE), or cotreatment including 100 ng/mL LPS and 100 nM of 16α-OHE, data was normalized by µg of protein in each sample (n=3-8 animals per group) **(B)**. Cell Supernatant was used to detect secreted ET-1 expression in control male and female *Bmpr2^+/R899X^* MT treated for 24 hours with a vehicle control, 100 ng/mL LPS, 100 nM 16α-OHE, or cotreated with 100 ng/mL LPS and 100 nM of 16α-OHE by dot blot analysis. *Inset:* representative dot blot image obtained using LI-COR Odyssey Imager. Data were normalized by protein concentration and cell number (n=3-5) **(C)**. The Shapiro-Wilk test was used to determine normality. Then, the data were analyzed using a two-way ANOVA followed by a Bonferroni *post hoc* test. *p<0.05, **p<0.001,***< 0.0001,****< 0.00001, ns, non-significant.

**Figure 7:**
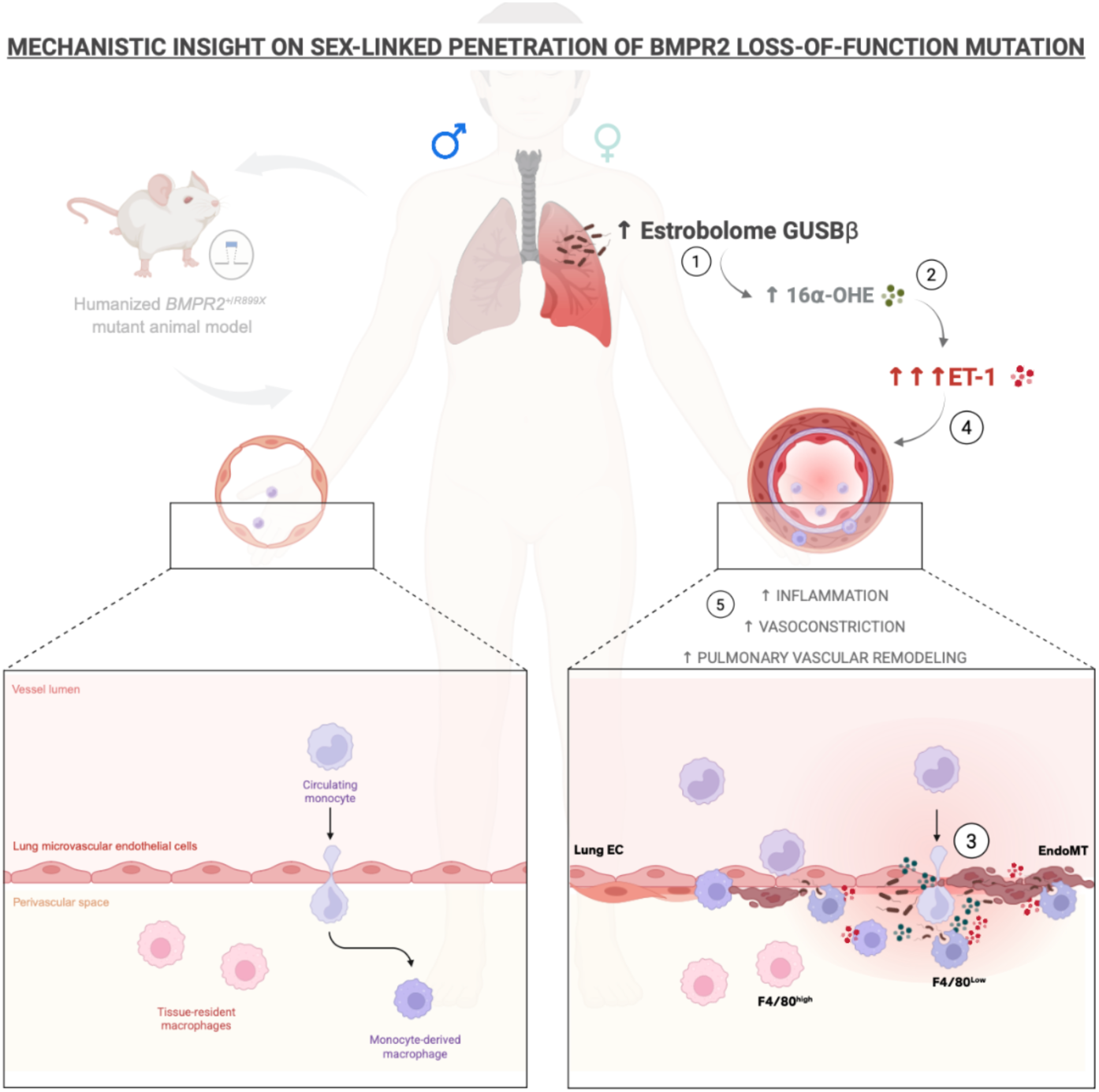
Mechanistic insight into sex-linked penetrance of *Bmpr2* loss-of-function mutation in pulmonary vascular disease. Schematic representation of a proposed mechanism linking sex-specific factors to enhanced disease penetrance in a female *Bmpr2* loss-of-function mutation carrier. A humanized *Bmpr2^+/R899X^* mutant animal model is depicted (left), illustrating the baseline vascular phenotype compared to the exacerbated remodeling observed in females. In females, alterations in the lung microbiome (estrobolome), characterized by increased β-glucuronidase (GUSB) expression and activity, promote enhanced deconjugation and availability of estrogen metabolites **(1)**. This shift favors the accumulation of the estrogen metabolite 16α-hydroxyestrone (16α-OHE), which has been associated with pathogenic signaling in the pulmonary vasculature **(2)**. Elevated 16α-OHE contributes to increased endothelin-1 (ET-1) production within the pulmonary microenvironment. This is accompanied by recruitment of circulating monocytes to the pulmonary microvasculature, their transmigration across the endothelial barrier, and differentiation into pro-inflammatory, monocyte-derived macrophages within the perivascular space **(3)**. The combined effect of increased ET-1 signaling and macrophage accumulation promotes endothelial dysfunction, inflammation, vasoconstriction, and progressive vascular remodeling, leading to luminal narrowing, increased vascular resistance, and pulmonary arterial hypertension (PAH) **(4)**. This model summarizes a novel putative sex-dependent axis involving the lung microbiome-16α-OHE-ET-1 pathway that amplifies immune-vascular crosstalk and promotes pulmonary vascular disease in PAH.

## IV. Discussion

Research on the lung microbiome remains a relatively new field, emerging over the past 15-20 years.^35^ Previously thought to be a sterile environment, it is now well-known that the lungs host a diverse population of microbial species^35^ that are essential for maintaining lung health, but also, when dysbiotic, are associated with a number of chronic lung diseases.^36^ Similar to findings in the MT females, studies using a Schistosomiasis-associated PH (Sch-PH) model reported reduced Ascomycota percentage,^19^ suggesting that a reduction in this phylum may play an important role in PH development. To the best of our knowledge, this work is the first to report evidence that a PAH-linked mutation may result in sex-linked lung microbiome dysbiosis.

Altered lung microbiome dynamics may contribute to the higher LPS concentration observed in the lungs of MT females. In the gut microbiome, it is well established that an altered composition can lead to increased LPS levels, resulting in chronic inflammation.^37^ This could begin to explain the sex-specific increased penetrance of the disease. Moreover, we reported a change in the lung estrobolome profile as a result of *Bmpr2* mutation, which could contribute to elevated levels of ꞵ-Glu and, subsequently, to the reabsorption of 16α-OHE.^24^ Although the difference in ꞵ-Glu in the whole lung tissue of MT males and females was not significant, ꞵ-Glu expression in the inner endothelial lining of the female microvasculature was higher than that of their male counterparts. To the best of our knowledge, this is a previously unnoted observation. The initial lack of statistical significance in the analysis of ꞵ-Glu by ELISA may be due to the preparation of the lung samples. It is important to note that these samples were homogenized using standard protease and phosphatase inhibitors specific to mammalian cells. It is possible that ꞵ-Glu produced by the estrobolome was denatured between the time of sample collection and the completion of the ELISA analysis. Future studies measuring ꞵ-Glu in the lungs should consider using both protease and phosphatase inhibitors suitable for both mammalian and microbial cell types, since ꞵ-Glu genes can be expressed by both. With the emergence of the microbiome area, it may be imperative to highlight this detail when assessing microbial protein mixed with host cells. Importantly, ꞵ-Glu is an enzyme that catalyzes the conjugation of estrogens into active forms, increasing levels and bioavailability of 16α-OHE. ^24,25^ These findings suggest that ꞵ-Glu may contribute to the elevated levels of 16α-OHE in MT females not because it is present in higher quantities throughout the lungs, but rather through spatially localized mechanisms in the endothelial lining of the lung microvasculature. Notably, this region of the lungs also exhibits an intense immune response, which may be explained by reabsorbed 16α-OHE, as evidenced by elevated ꞵ-Glu concentration and increased antigenic molecules.

Due to increased LPS in the lungs of MT females, we observed a severe infiltration of CD45+ macrophages that are capable of producing ET-1. This may explain why ET-1 levels are elevated in MT females with increased RVSP.^7^ Other studies demonstrate increased ET-1 in BMDM with *Bmpr2* mutations;^16^ however, to the best of our knowledge, we are the first to report that this response may be sex-specific. Additionally, 16α-OHE elevates ET-1 levels in MT females, suggesting that this may be one mechanism by which 16α-OHE contributes to the pathogenesis of PAH. As mentioned, ET-1 has been shown to play a role in the phenotypic transition of vascular endothelial cells to a myofibroblast-like phenotype via Endothelial-to-Mesenchymal Transition (EndoMT). Vascular cells that have undergone this process express significant levels of α-smooth muscle actin (α-SMA), a marker also observed in the microvasculature of mutants in this animal model,^7^ leading to elevated RVSP and progression of PAH This could begin to explain the estrogen paradox. Although estrogen has been described as cardioprotective in PAH,^13^ data suggest that the metabolite 16α-OHE may contribute to the increased prevalence observed in women.^27,38^ Notably, cotreatment with LPS and 16α-OHE was required to observe a significantly greater response in BMDM from females, indicating that both are required for the response observed in physiological conditions. Interestingly, 16α-OHE has been shown to promote local and systemic inflammation in other tissues.^39^ This data provides great insight into the complexity of PAH pathogenesis and paves the way for potential novel targets for the treatment of this severe disease.

This study strives to reveal previously unrecognized mechanisms of the pathogenesis of sex-linked PAH; however, several limitations should be acknowledged. The murine *Bmpr2* model is a valuable tool for gaining more insight into the pathogenesis of the disease, but it is important to recognize that mice and humans differ biologically. PAH is a complex disease that depends on multiple factors for development, and all murine models may have limitations.^40^ Additionally, there are also differences in the composition and diversity of the microbiota between humans and mice. Although murine models can be effective, our limited human tissue analysis may limit the translatability of these microbial differences to humans with this disease.^41^ Our data show a potential association between elevated 16α-OHE, ET-1, and ꞵ-Glu in females, as well as increased perivascular immune cell accumulation, with *Bmpr2^+/R899X^* mutation and PH. However, further studies are needed in order to establish causality. Another limitation is the lack of control of cycle variation, particularly in female samples. The fluctuation of sex hormones may influence estrobolome activity, estrogen metabolism, and ultimately the levels of 16α-OHE recorded. Despite these limitations, our findings provide important insight into the mechanisms of sex-linked PAH. These studies lay the groundwork for identifying causal relationships in the context of the disease and open the door to future, larger translational work.

## VI. Acknowledgment and Funding Support

We thank the Lab technician, MSc Maricela Castellon, for colony maintenance and sample collection; Dr. Ygor Marinho for partial microbiome-related analysis and lung sample preparation for MS/LC analysis; Dr. Nicholas W. Morrell (Professor of Medicine, University of Cambridge, United Kingdom) for generating and providing the *Bmpr2^+/R899X^* mutant mice and for exceptional consulting support, and; Dr. Amelia Bartholomew (UIC Department of Surgery) for facilitating collaborative flow cytometry studies. We also thank the UIC Research Resources Core (RRC), specifically the Mass Spectrometry Director, Dr. Hui Chen, for technical support with MS-LC measurements; Dr. Maria Sverdlov; and the Imaging Core Director, Ryan Deaton, for technical support with histological sectioning and slide scanning. This work has been supported by the National Institutes of Health (NIH HL159037), the UIC Dept. of Anesthesiology, and the UIC Health Equity Pilot Project Award (SDO). This project was partly supported by a 2022-23 UIC L@S GANAS Fellowship, a 2023 AHA ATVB Travel Grant, a 2023-2024 UIC Chancellor’s undergraduate research award (CURA) Scholarship, a 2025 Honors Research Grant Award, and a 2026 American Thoracic Society (ATS) Scholarship (selected as a late-breaking abstract and for student oral presentation (SPATS)).

## VI. Disclosures

No conflicts of interest declared by the authors.

## VII. Ethics Statement

All animal experiments in this study were performed as approved by the University of Illinois Chicago Institutional Animal Care and Use Committee.

## Notes

### Competing Interest Statement

The authors have declared no competing interest.

