## Supplementary material for "Sex-linked Lung Estrobolome May Contribute to Pulmonary Hypertension Penetrance of *Bmpr2 R899X* Mutation via an ET-1^high^ Endoregulatory Macrophage Phenotype": Suppl Material

### Supplemental Figures

### Suppl 1. Flow Cytometry Experimental Setup

A.

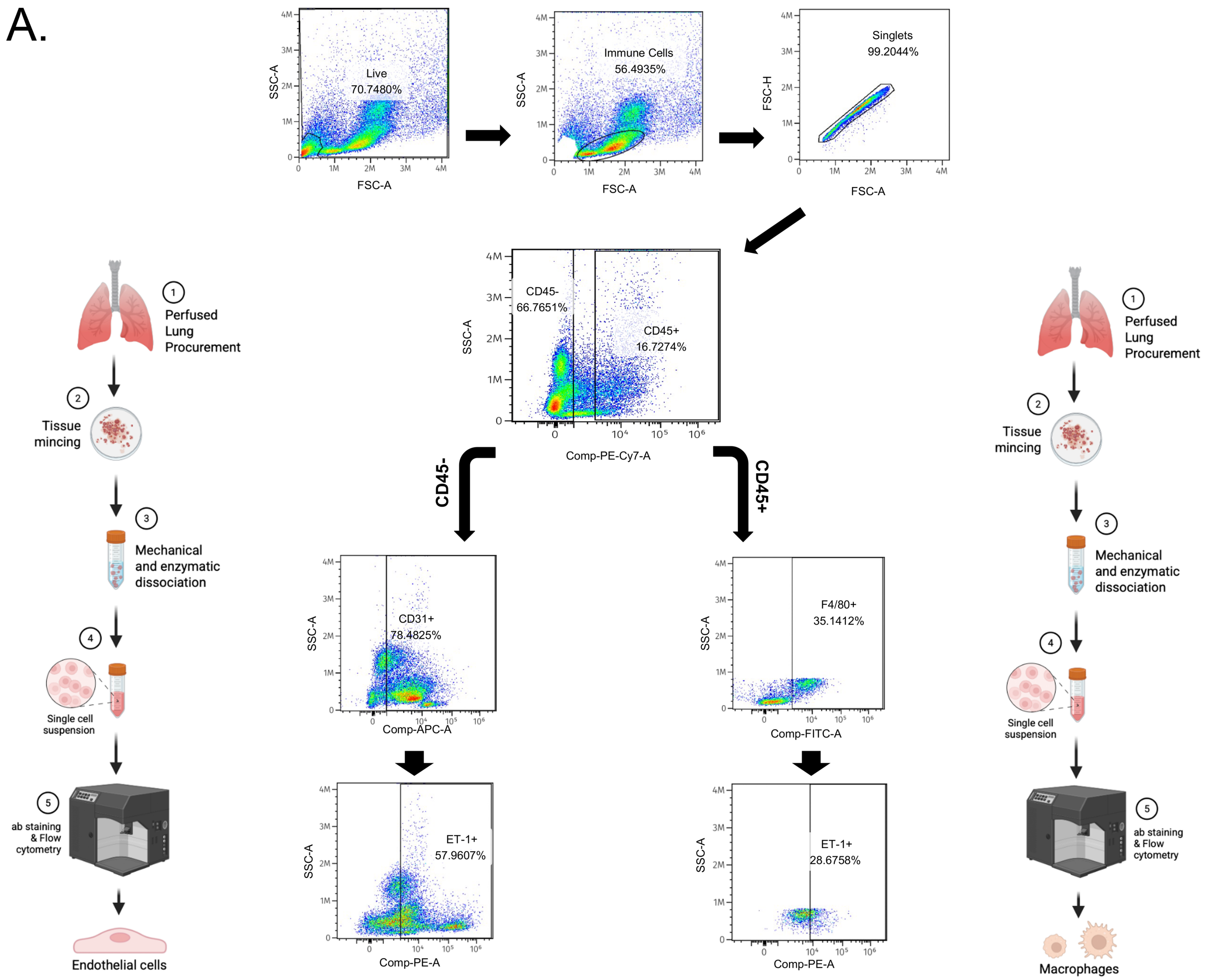


**Figure Suppl 1: Flow cytometry gating strategy.** Gates were selected to include CD45$^{-}$, CD31$^{+}$, ET-1$^{+}$ cells through Flow cytometry analysis to determine the percentage of endothelial cells containing ET-1 (Left). CD45$^{+}$, F4/80$^{+}$, ET-1$^{+}$ gates were used to analyze the percentage of macrophages expressing ET-1 (Right) **(A).** Flow Cytometry Analysis was done on 12-month-old male and female wildtype and *BMPR2$^{+/R899X}$* mutant animals. Flow cytometry analysis was also performed to measure the intensity of the cell markers used.

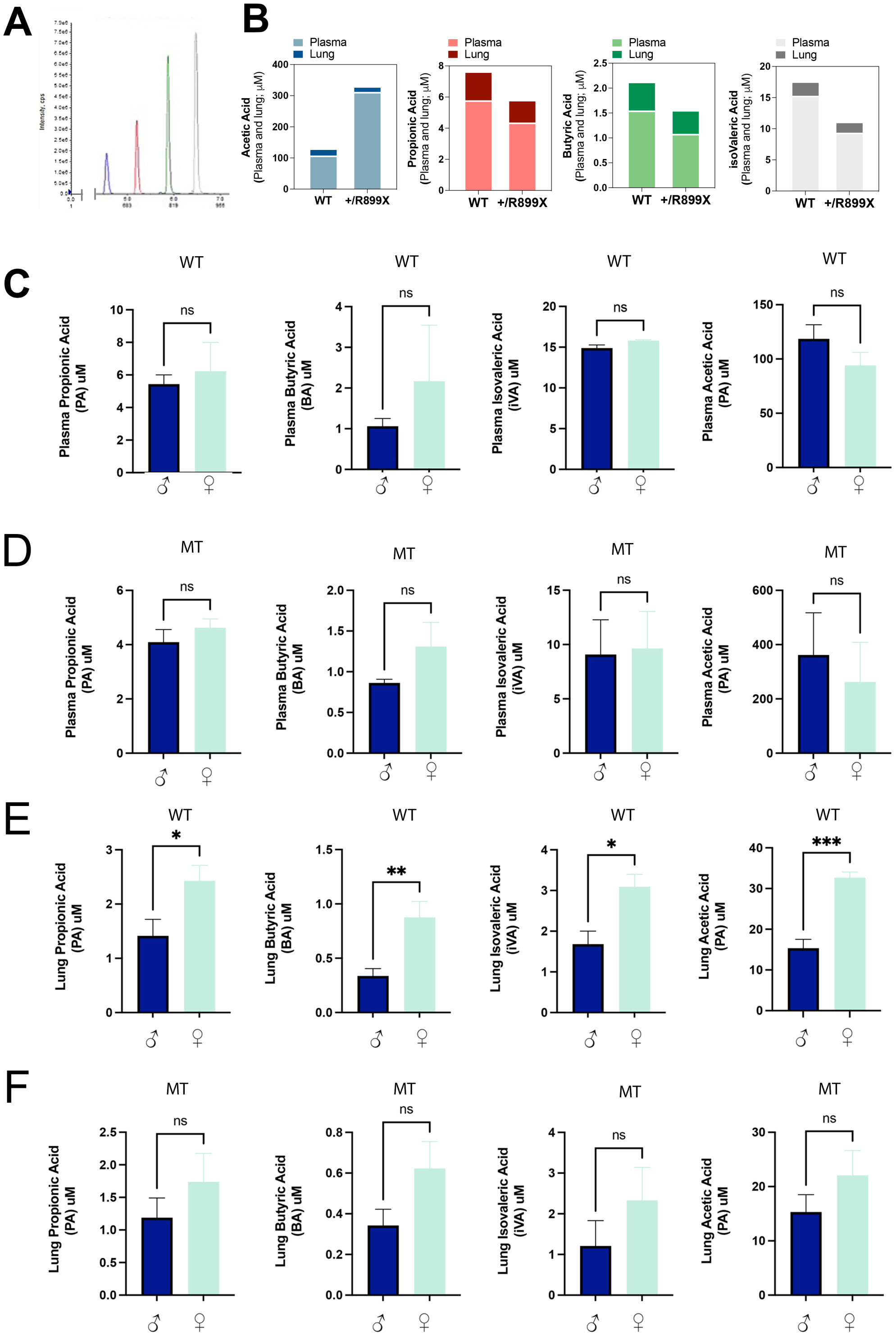


**Figure Suppl 2. Sex-linked short-chain fatty acids in humanized *BMPR2*$^{+/R899X}$ mutant mice model of pulmonary hypertension.** Lung tissue from 12-month-old male (dark blue) and female (light blue) wildtype control (WT) and *Bmpr2*$^{+/R899X}$ mutant mice (MT) were used for liquid chromatogram / MassSpec (LC/MS). LC/MS of the 4 short-chain fatty acids (SCFAs) analytes: acetic acid, propionic acid, butyric acid, and isovaleric acid measured in the plasma and cryopulverized lung samples from WT and *Bmpr2*$^{+/R899X}$. Normally distributed data are presented as the arithmetic mean +/- Standard Error of the Mean. The Shapiro-Wilk test was used to assess data normality, and the Brown-Forsythe or F test was used to assess variance equality. Then, a parametric (unpaired Student *t*-test) or nonparametric (Mann-Whitney test) test was performed accordingly. All tests were two-sided, and P < 0.05 was considered statistically significant. (n= 8-10 mice/group; ns = non-significant; *P < 0.05; ** P < 0.01; **** P < 0.0001).

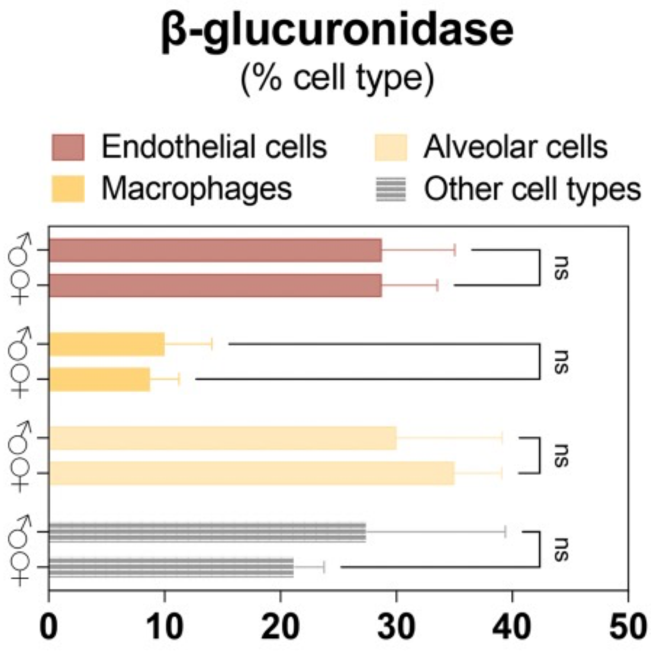


**Figure Suppl 3:** Quantification comparing the expression of β-Glu between male and female in various human cell types, such as Endothelial cells (Red), Macrophages (Orange), Alveolar (Yellow), and other (Grey). Analysis was done using the Human Protein Atlas (Karlsson et al., 2021. Sci Adv. 7(31): eabh2169); (proteinatlas.org).
